# Barrier mitigation measures trigger the rapid recovery of genetic connectivity in five freshwater fish species

**DOI:** 10.1101/2021.12.05.471259

**Authors:** Jérôme G. Prunier, Géraldine Loot, Charlotte Veyssiere, Nicolas Poulet, Simon Blanchet

## Abstract

Rivers are heavily fragmented by man-made instream barriers such as dams and weirs. This hyper-fragmentation is a major threat to freshwater biodiversity and restoration policies are now adopted worldwide to mitigate these impacts. However, there is surprisingly little feedback on the efficiency of barrier mitigation measures in restoring riverine connectivity, notably for non-migratory fish species. Here, we implemented a “before-after genetic monitoring” of the restoration of 11 weirs in France using a dedicated genetic index of fragmentation (the F_INDEX_), with a focus on five fish species from two genera. We found that most obstacles actually had a significant impact on connectivity before restoration, especially the highest and steepest ones, with an overall barrier effect of about 51% of the maximal theoretical impact. Most importantly, we demonstrated for the first time that mitigation measures such as dam removal or fish pass creation significantly and rapidly improved connectivity, with –for some barriers-a complete recovery of the genetic connectivity in less than twelve months. Our study provides a unique and strong proof-of-concept that barrier removal is an efficient strategy to restore riverine connectivity and that molecular tools can provide accurate measures of restoration efficiency within a few months.

**Graphical Abstract:** 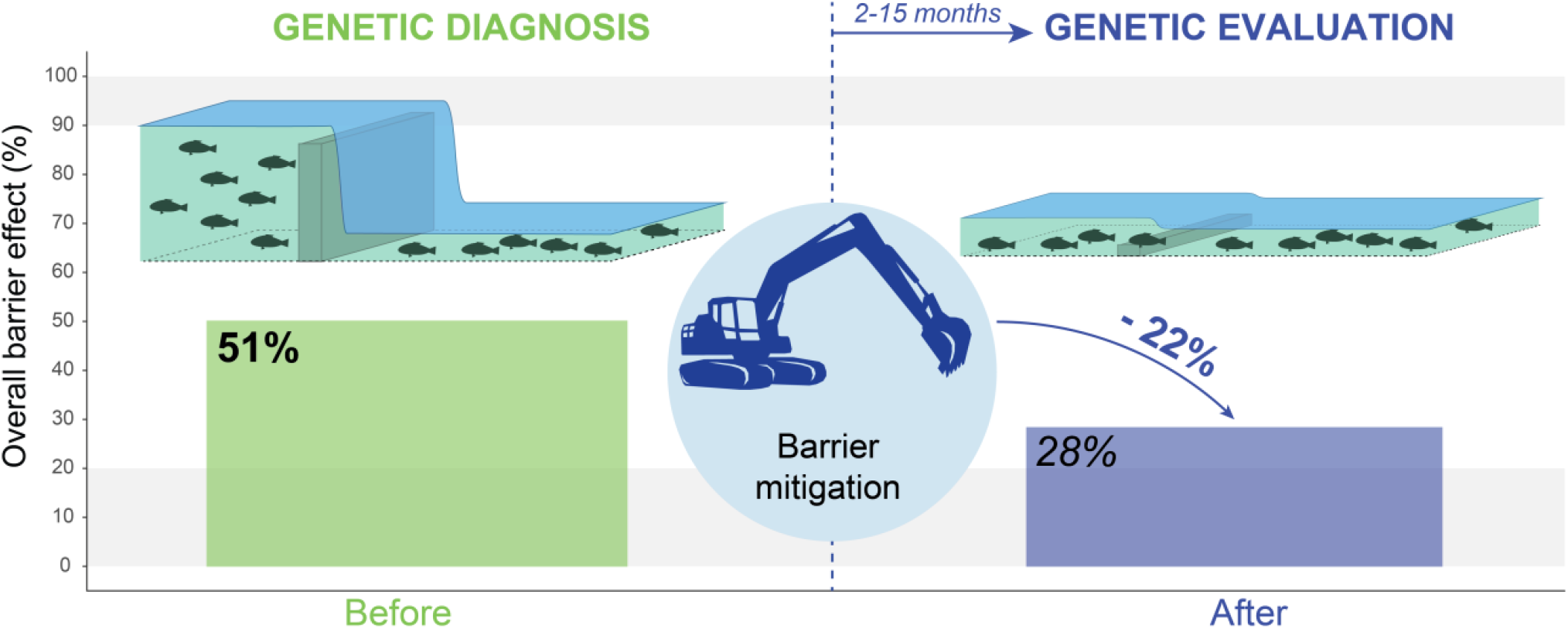

## 1. INTRODUCTION

Anthropogenic activities exert strong pressures on natural ecosystems, which alter both their physical and biological properties (Crutzen, 2006). This is especially the case for rivers that are highly fragmented by man-made instream barriers such as dams, weirs, water mills, etc. (Grill et al., 2019). In Europe for instance, more than one million obstacles have been reported (Belletti et al., 2020), representing 0,74 barriers per kilometer. Riverscape fragmentation affects the quality, the quantity and the accessibility of natural habitats, and thus prevents organisms to fulfill their life-cycle (Taylor et al., 1993). It is considered one of the most important threat to freshwater biodiversity (Sala, 2000). Given this “hyper-fragmentation” of rivers, the restoration of longitudinal (i.e., upstream-downstream) connectivity to promote fluxes of individuals and genes is hence considered a crucial step to recover the integrity of river ecosystems (Baguette et al., 2013; King & O’Hanley, 2016). Connectivity restoration is moreover the subject of restrictive legislations in many countries, such as in Europe with the Water Framework Directive (2000/60/EC), on which is based the EU’s biodiversity strategy for 2030 aiming at restoring at least 25,000 km of rivers to a free-flowing state (COM/2020/380).

The restoration of longitudinal connectivity implies barrier mitigation measures: the removal of obstacles, or, when removal is not an option (Blanchet & Tedesco, 2021; Lejon et al., 2009; Magilligan et al., 2017), the equipment of obstacles with natural or artificial fish passes (Seliger & Zeiringer, 2018; Silva et al., 2018). However, the actual efficiency of these barrier mitigation measures to restore genetic and/or demographic connectivity (Lowe & Allendorf, 2010) appear somehow unpredictable, depending on the type of river, the type of obstacle, the chosen type of restoration, as well as the timescale and the species considered (Rodeles et al., 2020). Dam removal has been found beneficial for the rapid recovery of some diadromous fish species (mainly salmonids) whose upstream migratory movements and thus spatial distribution were limited by the presence of barriers (Ding et al., 2019). However other organisms such as potamodromous fish (but also macroinvertebrates, macrophytes, etc.) may not (immediately) benefit from such removal (Brenkman et al., 2019; Gillette et al., 2016). Similarly, fish passes often show uneven levels of permeability across species, depending on pass design and maintenance as well as environmental conditions (Birnie-Gauvin et al., 2019; Harris et al., 2017; Noonan et al., 2012). These effects are yet still poorly documented, notably in the case of low-head structures (weirs, water-mills, sluices…) that do not benefit from the same attractivity as large dams in terms of public interest and research funding (O’Connor et al., 2015). Low-head structures are actually largely understudied when compared to their prevalence and the restoration efforts they represent (Belletti et al., 2020; Ryan Bellmore et al., 2017). Classical methods for the assessment of connectivity restoration efficiency, such as capture-mark-recapture, telemetry and monitoring of spatiotemporal changes in the composition of fish communities (Rodeles et al., 2020; Silva et al., 2018), cannot be systematically deployed at large management scales, especially for practitioners who are generally constrained by time and budget. For low-head barriers, mitigation measures are often undertaken opportunistically (Poff et al., 2003; Tonitto & Riha, 2016), with limited or coarse ecological monitoring of the system before restoration (Barry et al., 2018) and no (or limited) evaluation of the outcome in most cases (Cooke et al., 2019; Rodeles et al., 2017).

One challenge for practitioners is notably the lack of rapid and efficient connectivity assessment tools allowing both the *a priori* quantification of the individual impact of instream obstacles and the *a posteriori* quantification of the efficiency of implemented measures (removal or equipment; Cooke et al., 2019). If molecular tools are now commonly considered for the *a priori* assessment of barrier effects (Abernethy et al., 2013; Coleman et al., 2018; Dehais et al., 2010; Gouskov et al., 2016; Liu et al., 2020; Meldgaard et al., 2003; Prunier et al., 2018; Raeymaekers et al., 2009), there is still a surprising paucity of genetic studies dedicated to the temporal monitoring of changes in connectivity after restoration (Ding et al., 2019). Only a handful of recent studies could be identified, all focusing on the effect of the creation (Liu et al., 2020; Vega-Retter et al., 2020) or the removal (Fraik et al., 2021) of large dams on gene flow (but see Weigel et al., 2013). As a result, and despite the importance for practitioners and managers to assess (and communicate) the relevance of their actions for natural ecosystems, there are still very few indications that (i) current mitigation measures (barrier removal or fish pass creation) deployed worldwide are improving genetic connectivity (*sensu* Lowe & Allendorf, 2010) and (ii) that molecular approaches are efficient and operational tools to prioritize local restoration actions and assess their efficiency.

In this study, we implemented a “before-after genetic monitoring” of the restoration of 11 weirs in France using a recently developed genetic index of fragmentation (the F_INDEX_; Prunier et al., 2020). The F_INDEX_ provides, independently for each obstacle, an absolute and standardized estimate of (species-specific) genetic connectivity, while taking into account two confounding factors: the age of the obstacle and the size of populations on either side of the obstacle. Considering two common potamodromous fish genera, our objectives were (i) to quantify the initial impact of obstacles and thus to determine whether restoration was actually needed in the first place, and (ii) to quantify the gain in connectivity resulting from implemented restoration actions.

## 2. MATERIALS AND METHODS

### 2.1. Instream obstacles

The study took place at the scale of the French national hydrographic network. In close coordination with the French Office for Biodiversity (OFB) and French Water Agencies, we identified 11 obstacles (low-head dams <4m high) whose restoration was scheduled for the coming years (Figure 1; Table 1). These obstacles were located in the three largest French river basins (Garonne, Loire and Seine). On the basis of the consultation of old aerial photos and maps (French Ordnance Survey maps and cadastral plans), completed by surveys of local agencies, we estimated that they were constructed (or reconstructed after destruction or abandonment) between the 15^th^ and the 20^th^ century (Table 1). The obstacles ranged from 0.8 to 3.5m high and showed various slopes (Figure 1). Two of them were already equipped with a fish pass (Table 1). Each obstacle was described according to its height (< or ≥ 2m high) and slope (< or ≥ 45°) using a unique synthetic factor ‘Typology’ with four levels (‘low and gentle’, ‘low and steep’, ‘high and gentle’, ‘high and steep’). The restoration actions were conducted between 2015 and 2019 and in most cases (9 out of 11) consisted in the dismantlement of the obstacle. Note that we could not statistically assess the specific effects of fish passes on connectivity in the following analyses because there were too few of them, either before or after restoration.

**Figure 1:**
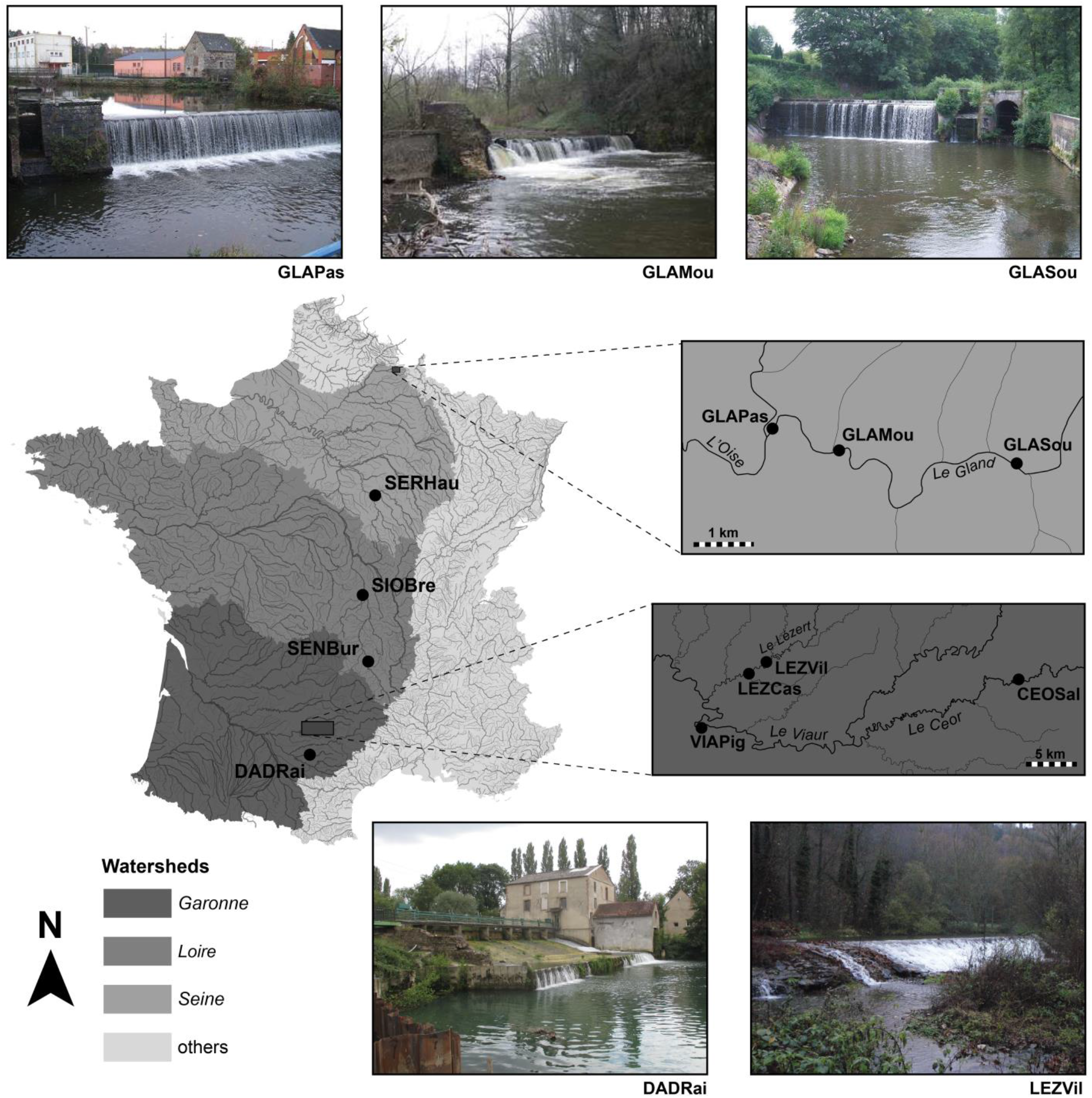
Geographic localization of studied instream barriers (black dots) in the main French watersheds.

**Table 1:** Main characteristics of obstacles (‘Lon’: longitude; ‘Lat’: latitude; ‘Height’ (in m), ‘Slope’ and presence of a ‘fish pass’), details about mitigation measures (type and date of actions) and timelag (in days) between mitigation measures and second sampling sessions.

| Obstacle characteristics |  |  |  |  | Mitigation measures |  |  |  |  |  |
| --- | --- | --- | --- | --- | --- | --- | --- | --- | --- | --- |
| Code | River | Lon | Lat | Creation | Height<br>(m) | Slope<br>(°) | Fish<br>pass | Action | Date | Timelag<br>(days) |
| CEOSal | Ceor | 2.57634 | 44.18213 | 1800 | 0.8 | > 45° | No | Removal | Jul-2016 | 427 |
| LEZCas | Lézert | 2.24649 | 44.18545 | 1400 | 1.1 | < 45° | No | Removal | Oct-2017 | 353 |
| GLAMou | Gland | 4.09026 | 49.92292 | 1800 | 1.2 | > 45° | No | Removal | Jul-2019 | 303 |
| VIAPig | Viaur | 2.18924 | 44.13735 | 1400 | 1.2 | > 45° | No | Removal | Aug-2017 | 44 |
| LEZVil | Lézert | 2.26765 | 44.19599 | 1800 | 1.9 | < 45° | Yes | Removal | Oct-2017 | 353 |
| DAD Rai | Dadou | 2.11982 | 43.78091 | 1800 | 2.0 | > 45° | No | Removal | Jun-2017 | 119 |
| SENBur | Senouire | 3.41640 | 45.27123 | 1500 | 2.2 | < 45° | No | Removal | Sept-2015 | 406 |
| GLAPas | Gland | 4.07951 | 49.92529 | 1800 | 2.6 | > 45° | No | Removal | Jul-2019 | 303 |
| SIOBre | Sioule | 3.29729 | 46.33353 | 1500 | 2.6 | < 45° | Yes | Fish pass restoration | Oct-2015 | 376 |
| GLASou | Gland | 4.11903 | 49.92129 | 1800 | 3.0 | > 45° | No | Removal | Nov-2016 | 344 |
| SERHau | Serein | 3.60304 | 47.92156 | 1830 | 3.5 | > 45° | No | Fish pass creation | Oct-2017 | 226 |

### 2.2. Biological models and Before-After sampling sessions

We focused on five common potamodromous species from two genera: minnows (*Phoxinus sp.: P. phoxinus* in the Seine*, P. fayollarum* in the Loire and *P. dragarum* in the Garonne watershed) and gudgeons (*Gobio sp*: *G. gobio* in the Loire and the Seine and *G. occitaniae* in the Garonne watershed). Within each genus, species are allopatric (Denys et al., 2020) but were here considered to have very similar life history traits and movement behaviors (Keith et al., 2011). These species are small insectivorous cyprinids (maximal body length of 140 and 200mm, in minnows and gudgeons respectively) that show distinct foraging strategies: minnows preferentially feeds in the water column, whereas gudgeons feed on the bottom (Keith et al., 2011). We selected these species because they are widespread and abundant and hence easy to catch for practitioners, which make them ideal models for an operational tool such as the F_INDEX_.

Sampling operations ‘Before’ and ‘After’ restoration were performed using electrofishing, until a maximum of 30 adult individuals of each species were captured on either side of obstacles. Fish were captured in the direct downstream and upstream vicinity of each obstacle (Prunier et al., 2020), starting from the downstream site to avoid accidental upstream to downstream movements of individuals. A piece of pelvic fin was sampled on all individuals and stored in 96% alcohol. All fish were returned alive to their sampling site. Electrofishing and fin sampling was performed according to legal authorizations and permits.

Both genera could be sampled at all sites except in DADRai (gudgeons only) and GLASou (minnows only). The sampling sessions after the restoration occurred on average 9.7 (± 3.9 SD) months after the end of the restoration (from 44 days in VIAPig to 427 days in CEOSal; Table 1). These after-restoration timelags could not be homogenized better, owing to logistic and administrative difficulties (e.g., high water levels preventing safe fieldwork).

### 2.3. Genotyping and F_INDEX_ computation

We considered 19 and 15 microsatellite markers in minnows and gudgeons, respectively. DNA extraction, genotyping and assessment of null alleles and gametic disequilibrium followed previously published procedures (Prunier et al., 2018, 2020; Appendix S1). For each dataset (combination of one obstacle and one genus; n = 20 since only one genus could be sampled at GLASou and DADRai; Table 2) and each passage (‘Before’ and ‘After’), we computed the F_INDEX_ and the standard deviation of the F_INDEX_ as detailed in Prunier et al. (2020) using a dedicated R-based pipeline. Briefly, the F_INDEX_ corresponds to the rescaling of pairwise measures of genetic differentiation within their theoretical range of variation given effective population sizes (approximated from expected heterozygosity) and the number of generations elapsed since barrier creation. The F_INDEX_ is therefore a standardized (across barriers and species) index expressed as a percentage, with values lower than 20% representing fully permeable structures and values higher than 90% representing total barriers to gene flow (see Prunier et al., 2020 for details). F_INDEX_ values are computed along with a standard deviation SD_F_ (or a 95% confidence interval CI_95%_) that takes into account biological uncertainty stemming from n_F_ = 4 parameters (two mutation rates and two metrics of genetic differentiation; Prunier et al., 2020). We considered a generation time of 2 years in minnows and 2.5 years in gudgeons to compute the number of generations elapsed since barrier creation (Kottelat & Freyhof, 2007).

**Table 2:** For each dataset (that is, a combination of one obstacle and one genus), observed effect sizes of barrier effects (F±SD_F_) both before and after restoration and observed effect sizes of restoration (ΔF± SE_Δ_), respectively. Note that SD_F_ = 2 x SE_F_. Significant effect sizes (see Figure 2) are in bold and indicated with a star. The last column ‘Recovery’ indicates whether the restoration action (Removal or Fish pass creation) led to the partial or the full recovery of connectivity, when applicable.

| Obstacle code | Genus | Barrier effect |  |  |  | Restoration efficiency |  |  |  |
| --- | --- | --- | --- | --- | --- | --- | --- | --- | --- |
| | | Before restoration | | After restoration | | Action | $\Delta F$ | $SE_\Delta$ | Recovery |
| | | F | $SD_F$ | F | $SD_F$ | | | | |
| CEOSal | Gudgeons | 0.00 | 0.00 | 0.00 | 0.00 | Removal | 0.00 | 0.00 | / |
|  | **Minnows** | **64.42** | **1.59** | * **35.52** | **2.22** | * Removal | **-28.91** | **1.37** | * **Partial** |
| LEZCas | **Gudgeons** | **27.25** | **6.26** | * 0.00 | 0.00 | Removal | **-27.25** | **3.13** | * **Full** |
|  | Minnows | 0.00 | 0.00 | 11.66 | 12.47 | Removal | 11.66 | 6.23 | / |
| GLAMou | Gudgeons | 0.00 | 0.00 | 0.00 | 0.00 | Removal | 0.00 | 0.00 | / |
|  | Minnows | 4.27 | 4.57 | 0.00 | 0.00 | Removal | -4.27 | 2.28 | / |
| VIAPig | **Gudgeons** | **32.48** | **0.99** | * 12.41 | 3.70 | Removal | **-20.07** | **1.91** | * **Full** |
|  | **Minnows** | **29.80** | **4.59** | * 8.35 | 8.93 | Removal | **-21.45** | **5.02** | * **Full** |
| LEZVil | Gudgeons | 0.00 | 0.00 | 10.39 | 11.14 | Removal | 10.39 | 5.57 | / |
|  | Minnows | 2.32 | 2.63 | 0.00 | 0.00 | Removal | -2.32 | 1.31 | / |
| DAD Rai | **Gudgeons** | **68.09** | **1.58** | * **26.82** | **1.20** | * Removal | **-41.27** | **0.99** | * **Partial** |
| SENBur | **Gudgeons** | **54.75** | **1.02** | * **48.22** | **1.87** | * Removal | **-6.53** | **1.07** | * **Partial** |
|  | Minnows | 14.59 | 15.60 | 0.00 | 0.00 | Removal | **-14.59** | **7.80** | * / |
| GLAPas | **Gudgeons** | **55.42** | **2.24** | * 0.00 | 0.00 | Removal | **-55.42** | **1.12** | * **Full** |
|  | **Minnows** | **42.88** | **2.87** | * 0.00 | 0.00 | Removal | **-42.88** | **1.43** | * **Full** |
| SIOBre | Gudgeons | 0.00 | 0.00 | 0.00 | 0.00 | Fish pass | 0.00 | 0.00 | / |
|  | **Minnows** | **51.84** | **2.57** | * **39.46** | **1.61** | * Fish pass | **-12.38** | **1.52** | * **Partial** |
| GLASou | **Minnows** | **49.26** | **0.76** | * **32.51** | **4.77** | * Removal | **-16.75** | **2.42** | * **Partial** |
| SERHau | Gudgeons | 0.00 | 0.00 | 0.00 | 0.00 | Fish pass | 0.00 | 0.00 | / |
|  | **Minnows** | **57.33** | **1.31** | * **49.10** | **2.05** | * Fish pass | **-8.23** | **1.22** | * **Partial** |

### 2.4. Barrier effects and restoration efficiency

We adopted a meta-analytical approach (Borenstein, 2009), considering each dataset (combination of one obstacle and one genus; n = 20) as an independent study providing two effect sizes (F_INDEX_ values before and after restoration ± SD_F_). Six datasets were non-informative (F_INDEX_ = 0 and SD_F_ = 0 before restoration) and were thus discarded, resulting in 14 datasets, of which three were associated with a non-significant barrier effect (see results). The interpretation of the F_INDEX_ being meaningful (barrier effect expressed as a percentage of maximum fragmentation), the F_INDEX_ value before restoration was directly used as the observed effect size F±SE_F_ of barrier effect in each dataset, with 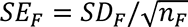. We used the raw difference ΔF(±SE_Δ_) between F_INDEX_ values computed after (‘treatment’) and before (‘control’) restoration as the observed effect size of restoration for each dataset (Equations 4.2 and 4.6 in Borenstein, 2009). The raw difference ΔF can be directly interpreted as the (positive or negative) change in the amount of fragmentation following restoration and was thus preferred over the standardized mean difference Hedges’ g (Borenstein, 2009).

We used random-effect meta-regressions with moderators (metafor::rma.mv R-function; Harrer et al., 2021; Viechtbauer, 2010) to compute the overall true effect sizes of the barrier effects 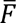 and of the restoration of obstacles 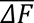, while taking into account different possible sources of variation in effect sizes: within-datasets (i.e., SE_F_ or SE_Δ_), between-datasets and, possibly, across different covariate modalities. Considered covariates were ‘Genus’ (two levels: minnows or gudgeons), ‘Typology’ (four levels: see above), as well as, in the case of 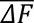, the ‘timelag’ (two levels: < 1 year or > 1 year) between the restoration operation and the second sampling session. To account for within-dataset variability (SE_F_ or SE_Δ_), we used dataset ID as an outer random grouping factor. To allow residual heterogeneity to differ across covariate modalities, each covariate was successively defined both as a moderator and an inner random grouping factor. Models were run with a diagonal variance-covariance matrix as a random effect structure. Covariates identified as significant moderators were then kept as inner random grouping factors in final models without moderator to get final estimates of the true effect sizes 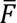 and 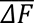 along with their respective 95% confidence interval CI_95%_.

## 3. RESULTS

Altogether, a total of 1049 and 1038 genotypes could be obtained from the various sampling sessions (through space and time) in minnows and in gudgeons, respectively, with an overall mean of 26.1 (± 3.2 SD) genotypes per genus, sampling site and sampling session. The mean number of loci across datasets was 15.7 (± 1.8 SD) and 14.4 (± 0.8 SD) in minnows and in gudgeons, respectively.

F_INDEX_ values ranged from 0 to 68.1 % (mean: 27.7 ± 25.8 SD) before restoration, with a significant barrier effect detected in 11 out of 20 datasets (Table 2). Genera showed highly contrasted responses to obstacles: all obstacles but GLAMou and LEZVil had a significant impact on connectivity before restoration, but only VIAPig and GLAPas did impact both genera simultaneously (Table 2; Figure 2). Accordingly, ‘Genus’ was not identified as a significant moderator of the overall effect size 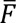 (Table 3). Only ‘Typology’ did significantly influence 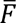, with steep obstacles higher than 2 m showing an overall effect size 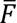 40 % (CI_95%_ = [14.2; 65.8]) higher than gentle weirs lower than 2 m (Table 3; Figure 3). Once all possible sources of variation in effect sizes were taken into account (both within-datasets and between modalities of typology), the final true effect size of fragmentation 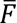 was of 51.4 %, a value significantly different from 0 and higher than the F_INDEX_ significance threshold of 20% (CI_95%_ = [45.1; 57.7]).

**Figure 2:**
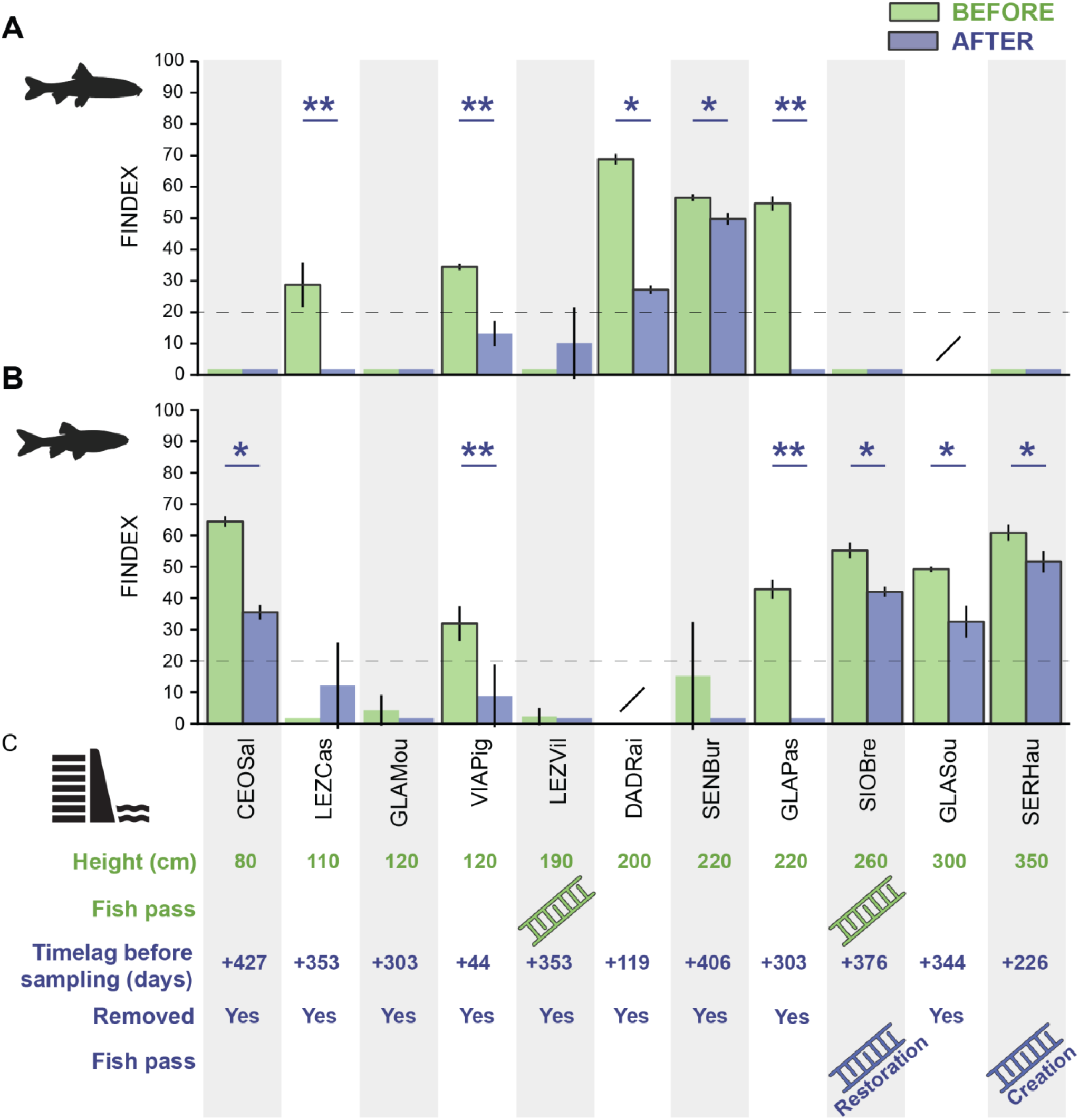
Main results of the before-after genetic monitoring. For each obstacle (in columns), bars represent F_INDEX_ values with 95% confidence intervals as computed before (in green) and after (in purple) restoration in gudgeons (panel A) and minnows (panel B). Slashes indicate no data in both A and B. Outlined bars represent significant barrier effects (F_INDEX_ > 20%). Green stars indicate a significant change in F_INDEX_ values after restoration (non-overlapping confidence intervals). Double purple stars indicate the full recovery of connectivity following restoration (see details in Table 2). Panel C also provides few details about obstacles (in green) and restoration (in purple) for direct comparisons with F_INDEX_ values (see Table 1). Obstacles are sorted by their increasing height. Note that F_INDEX_ values actually increased after restoration in two datasets (LEZVil in gudgeons and LEZCas in minnows), but that new values were lower than 20 % and non-significantly different from 0 (and thus from F_INDEX_ values before restoration) according to CI_95%_, a phenomenon owing to the stochasticity of genetic approaches

**Figure 3:**
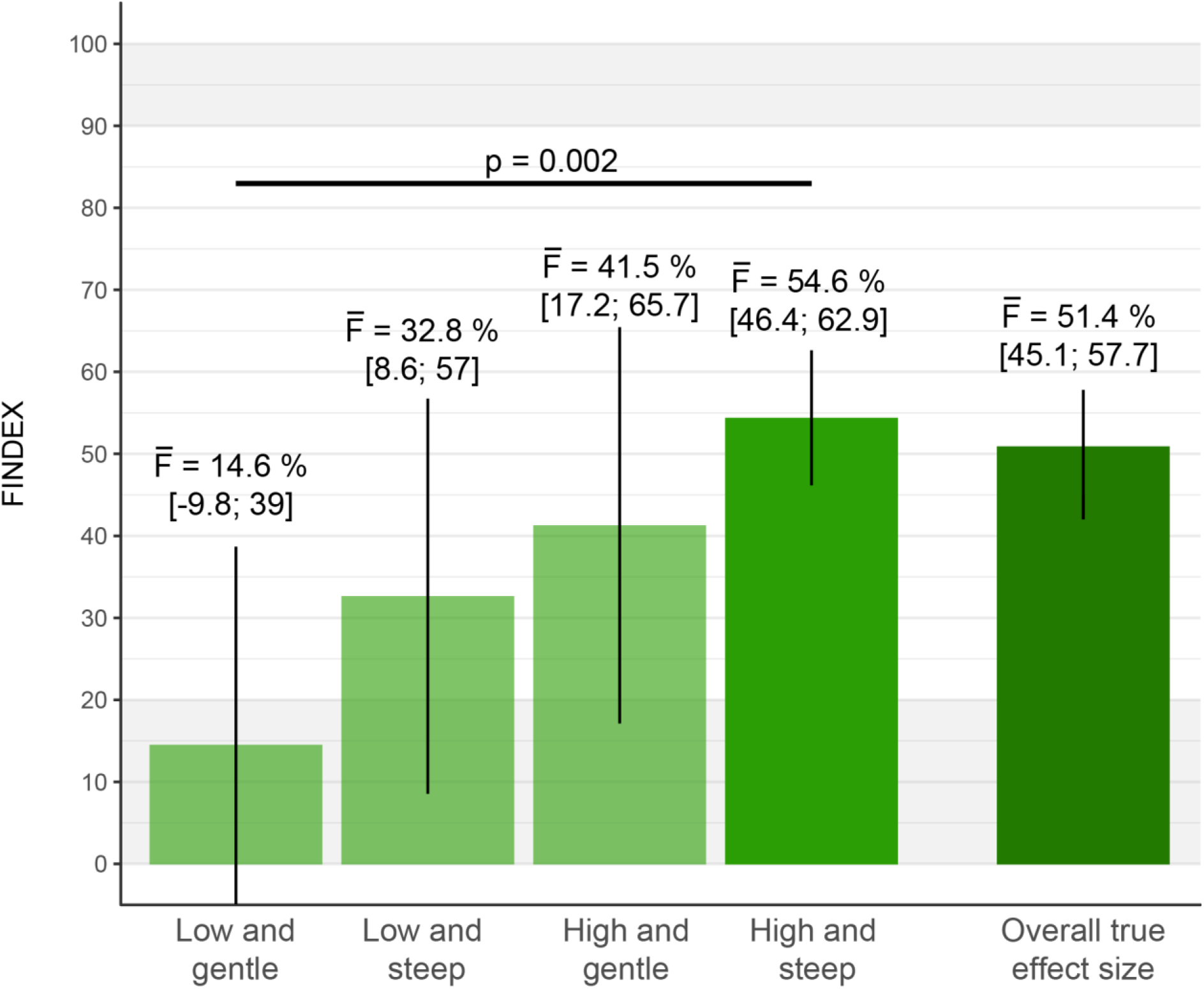
Overall effect sizes 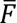 of fragmentation for each modality of typology and overall true effect size 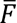 taking all significant sources of variation (within datasets and across modalities) into account. Washed out bars indicate that the effect size of fragmentation is not significant (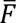 > 20% or CI_95%_ including 20%; Prunier et al., 2020).

**Table 3:**
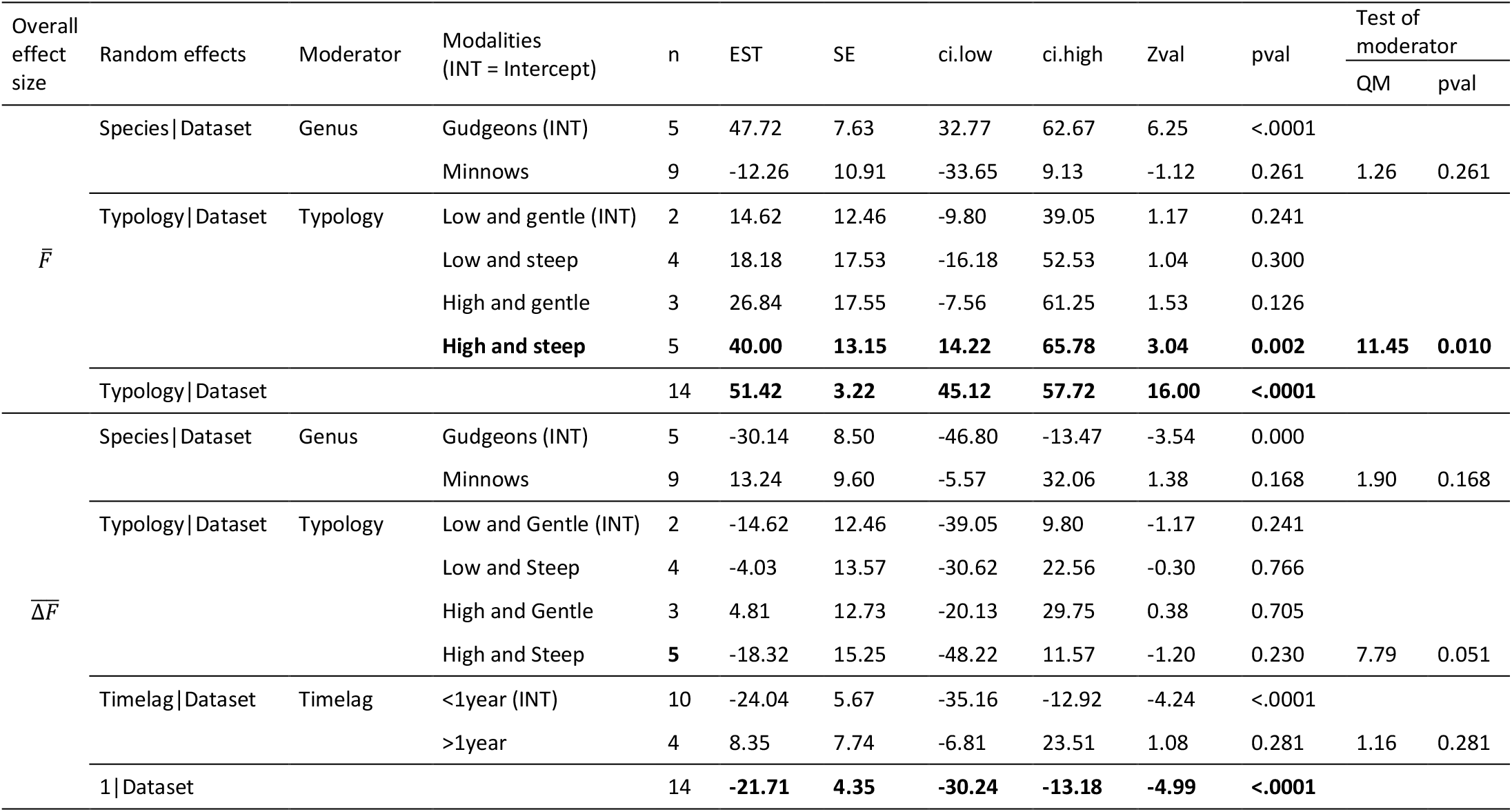
Results of random meta-analyses for the overall effect sizes of fragmentation 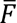 and of restoration 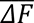 In presence of a moderator, EST is the estimate of the overall effect size for the intercept (INT) and the deviation from the intercept for the alternate modalities, with the moderator as a random effect. In absence of moderator (indicated with a slash), EST is the final estimate of the overall effect size taking into account all sources of variation (including the previously identified significant moderators as random effects, when applicable). Also provided are the number *n* of datasets in each modality, the standard error (SE) and CI_95%_ (ci.low and ci.high) around EST, the Wald-Type Z statistic (Zval) and the associated p-value (pval), and the QM test of moderator effect (QM statistic and associated p-value).

Except when obstacles had no effect before restoration (F_INDEX_ < 20%), in which case restoration had no effect either (F_INDEX_ < 20% after restoration; 5/10 obstacles in gudgeons and 4/10 obstacles in minnows), restoration systematically led to a significant decrease in F_INDEX_ values (as indicated by non-overlapping confidence intervals; Figure 2), with F_INDEX_ values after restoration ranging from 0 to 49.1 % (mean: 13.7 ± 17.8 SD). This decrease led to the full recovery of connectivity (F_INDEX_ < 20% after restoration) in 3/5 obstacles in gudgeons and 2/6 obstacles in minnows. The observed effect sizes of restoration ΔF ranged from -55.4% to +11.7% (mean: -14 ± 18 SD; Table 2). Neither ‘Genus’, ‘Typology’ or ‘Timelag’ were identified as significant moderators of the overall effect size of restoration 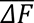 (Table 3, Appendix S2). The final overall effect size of restoration 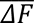 was of -21.7%, a value significantly different from 0 (CI_95%_ = [-30.2; -13.2]). Overall, the effect of restoration was thus the same across genera, did not depend on the typology of the obstacle and, interestingly, was independent from the timelag between the restoration and the second sampling session. In other words, we found evidence that genetic connectivity could be recovered (entirely in some cases, partly in most cases) in just a few months after restoration.

## 4. DISCUSSION

Quantifying the impact of instream barriers on potamodromous fish species as well as the efficiency of mitigation measures is primordial in the context of restoration planning, so as to properly allocate limited resources towards the most impactful obstacles, inform trade-offs between ecological and socio-economic issues, and refine restoration techniques (Hermoso et al., 2012; Rodeles et al., 2020; Silva et al., 2018). Quantification is yet a difficult task, notably because of technical and financial constraints preventing the parallel monitoring of multiple obstacles and because of the relative lack of operational tools allowing valid comparisons across both contexts and species (Cayuela et al., 2018). The response of freshwater organisms to connectivity restoration has often been studied at the community- or at the population-levels (Brenkman et al., 2019; Frey, 2021; Magilligan et al., 2021; Muha et al., 2021; Stanley et al., 2002; Sun et al., 2021), rarely at the genetic level (Fraik et al., 2021), and our study is the first to document the systematic and rapid recovery of gene flow over a series of independent restoration actions. In this study, we used a standardized genetic index of fragmentation to quantify both the impact of 11 low-head dams on gene flow in five freshwater fish species, and the efficiency of mitigation actions in restoring genetic connectivity.

Before restoration, we found a significant barrier effect in 11 out of 20 datasets, with two obstacles showing no impact on any genus and two obstacles significantly impacting both genera. Surprisingly, out of nine obstacles with genetic data in both genera, five showed large discrepancies in genus response to fragmentation, with either only gudgeons or only minnows significantly impacted (Figure 2). These discrepancies illustrate how barrier effects can be highly species- or genus-dependent (Amaral et al., 2021; Blanchet et al., 2010; Prunier et al., 2018), and thus hardly predictable given our limited knowledge about fish movement behavior and capacities (Baudoin et al., 2014; Thurow, 2016). In absence of a dedicated fish pass, individuals are supposed to take advantage of drowned conditions, that is, of periods where water level rises above the height of the dam, to cross the obstacle (Keller et al., 2012). However, such propitious conditions of obstacle drowning might not be encountered every year, at all localities, and equally across all species/individuals, depending on their swimming behavior and capabilities in various environmental conditions and to the timing of submersion compared with the timing of individual movements (Carpenter-Bundhoo et al., 2020; Holthe et al., 2005; Keller et al., 2012).

This may explain why F_INDEX_ values differed so much across datasets, and why the overall effect size of fragmentation 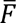 was unrelated to the considered genus. However, 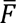 was significantly influenced by the typology of obstacles, with high and steep obstacles showing an overall effect size 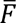 40% higher than low and gentle obstacles (Table 3, Figure 3). This finding is in line with classical expectations about the impact of dam typology (e.g., Januchowski-Hartley et al., 2019) and conclusions from other studies (e.g., Amaral et al., 2019; Keller et al., 2012; Zigler et al., 2004), as well as with the obstacle drowning hypothesis: the highest dams (≥ 2 m) might rarely be drowned (leading to the increase in 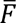), but, in presence of a gentle slope (< 45°), they might become partly crossable by some individuals, at least at intermediate drowning conditions, so that only the highest and steepest obstacles have an overall significant effect size (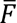 = 54.6%; Figure 3). This result should of course be confirmed and refined with additional datasets, but it provides a relevant and meaningful benchmark for practitioners to adjust restoration planning even in the absence of any individual quantification of barrier effects.

With this effect of typology taken into account, we estimated the true effect size of fragmentation as of 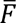 = 51.4 %: in other words, we might expect a 51 % decrease in gene flow in presence of a low-head dam (irrespective of the species or the context). Given these different findings, it appears that a preliminary diagnosis based on a standardized genetic tool such as the F_INDEX_ may actually help managers quantify and compare barrier effects across species and obstacles, and thus orientate their restoration efforts towards the most problematic structures (Prunier et al., 2020), keeping in mind that the most impactful obstacles might be the highest and steepest ones, but that more seemingly anodyne obstacles might also be particularly impacting for some species. Of course, we willingly acknowledge that other ecological and socio-economic indicators should be considered in restoration planning as well (Hermoso et al., 2012). It is also noteworthy that the F_INDEX_ might help evaluate the species-specific efficiency of fish passes, an important step to drive future technical developments (Foulds & Lucas, 2013): the two considered fish passes before restoration appeared beneficial in all situations except for minnows at SIOBre (Figure 2), illustrating the challenge of locally designing passes adapted to different fish species with distinct life-history traits and requirements (Birnie-Gauvin et al., 2019; Silva et al., 2018).

Our most striking result concerns the efficiency of mitigation measures. All restoration actions led to a significant reduction of barrier effects, provided there was an actual barrier effect in the first place. We quantified an overall 22% decrease in fragmentation levels following restoration. This means that mitigation measures may allow the full recovery of genetic connectivity for any initial fragmentation level of up to 42% (i.e., 20% < F_INDEX_ ≤ 42%), a value to be compared to the overall effect size of fragmentation 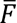 = 51% before restoration. Most interestingly, this systematic gain in connectivity was achieved within a few months after restoration only. For instance, weir removal led to the full recovery of genetic connectivity at three localities (LEZCas, VIAPig and GLAPas), indicating that it ensured the full mixing of individuals, and thus of allelic frequencies, within a year, and even within two months only at VIAPig. However, not all restoration actions proved equally efficient within the same timeframe, the recovery of connectivity being only partial in several situations. This is the case of the two (new or improved) fish passes at SERHau and SIOBre that resulted in an 8 to 12% recovery in genetic connectivity in minnows within a year. These reductions are highly encouraging, but it is still unknown whether recovery is still ongoing: further temporal monitoring is needed to detect migration-drift equilibrium and determine the final gain in connectivity following restoration. Furthermore, and although weir removal is expected to be more efficient than fish pass creation as a restoration option (Birnie-Gauvin et al., 2019; Sun et al., 2021), other removal actions only led to a partial recovery of connectivity, sometimes even after a year (e.g., at CEOSal in minnows). The migration behavior of fish being still poorly documented and likely to deeply differ across species and contexts, it probably explains why the true effect size of restoration was unrelated to the considered genus, the typology of the obstacle or the timelag between restoration and sampling. We can only speculate on why close-range genetic mixing would sometimes take so long despite the absence of any obstacle to movement. For instance, weir removal might result in profound changes in upstream and downstream habitat characteristics (Bednarek, 2001; Doyle et al., 2005), locally inducing a temporary repelling effect on fish. It has also been suggested that, in some conditions, fragmentation might lead individuals to adjust their life-history strategies towards residency (Branco et al., 2017), which might further delay genetic connectivity after removal. Nevertheless, we expect the full recovery of genetic connectivity in the coming years, which will however require further genetic monitoring of these situations.

## 5. CONCLUSION

Our study provides a strong proof-of-concept that barrier removal and, probably to a lesser extent, fish pass creation, are efficient mitigation strategies to restore riverine genetic connectivity in just a few months. We also illustrated how before-after genetic monitoring based on a standardized tool such as the F_INDEX_ constitute a promising support for practitioners in the planning and the monitoring of restoration. We believe that the large-scale deployment of this methodology in the future, with a growing number of case studies, will make it possible to lift the veil on the complex links between individual crossing success, life history traits of organisms and barrier typologies.

## ACKNOWLEDGEMENTS

This study was financially supported by the Office Français pour la Biodiversité and the Région Occitanie. We warmly thank all the members of local agencies (Office Français pour la Biodiversité, Fédérations de Pêche, Syndicats Mixtes, Agences de l’Eau) that have been involved in this project and directly helped us fulfill our objectives, either by performing the sampling or by providing valuable information about local restoration actions.

## AUTHORS’CONTRIBUTIONS

JGP, SB and NP conceived the ideas and designed methodology; JGP and SB collected the samples in the field; CV and GL produced molecular data; JGP analyzed the data; JGP and SB led the writing of the manuscript. All authors contributed critically to the drafts and gave final approval for publication.

## APPENDIX S1: DNA EXTRACTION AND GENOTYPING

Genomic DNA was extracted using a salt-extraction protocol (Aljanabi & Martinez, 1997). A subset of 15 and 19 autosomal microsatellite loci from Grenier et al. (2013) were amplified and genotyped in gudgeons (BL1-153, Gob15, Gob16, Gob22, LC293, Lco4, MFW1, Ca1, CypG24, Gob12, Gob28, Lsou5, Rvla21177, Smv03 and Lro12) and minnows (BL1-153, Ca3, CtoA-247, CtoG-075, CypG9, LSou8, LleA-071, LleB-072, Ppro-132, Rru4, BL1-44, BL1-84, BL1-98, LC27, LceC1, LleC-090, Lsou5, MFW1 and Rhca20), respectively, following PCR conditions described in Grenier et al. (2013).

Each combination of a genus, an obstacle and a sampling session (n = 40) was considered a unique genotypic dataset, with corresponding genotypes coded in the *genepop* format (Rousset, 2008). For each dataset, we assessed the presence of null alleles and checked for gametic disequilibrium using the *null.all* function (R-package *PopGenReport*; Adamack & Gruber, 2014) and the *test_LD* function (R-package *genepop*; Rousset, 2008), respectively. Any locus showing significant gametic disequilibrium and/or evidence of null alleles was discarded before F_INDEX_ calculation (see Table 2 in main text for outputs).

## APPENDIX S2: THE EFFECT OF TIMELAG ON THE OBSERVED EFFECT SIZES OF RESTORATION ΔF

The observed effect sizes of restoration ΔF (±CI_95%_) did not show any significant trend when increasing the timelag between restoration and sampling.

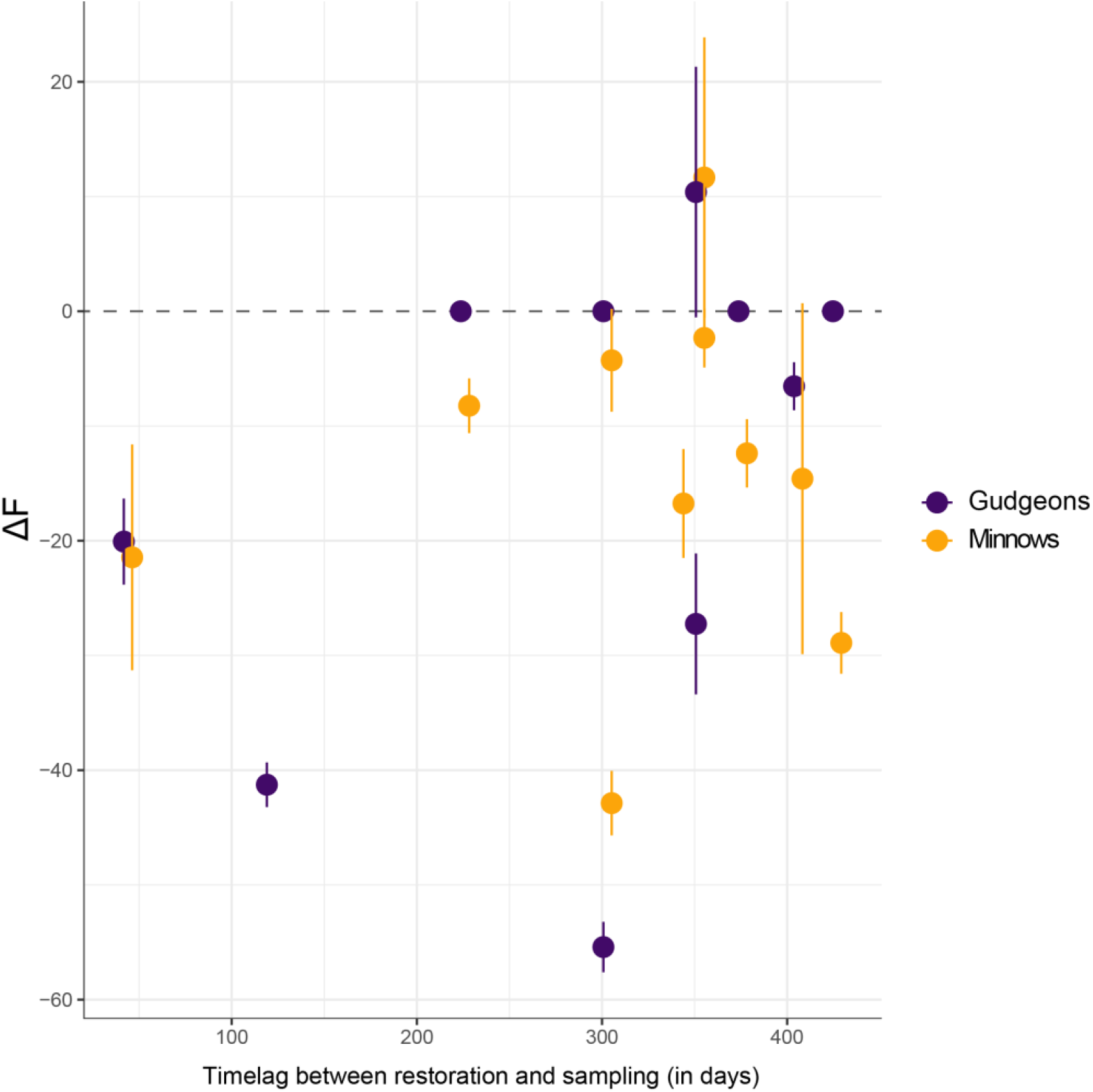

## Notes

### Competing Interest Statement

The authors have declared no competing interest.

